# Cutaneous human papillomavirus E6 impairs the cGAS-STING pathway

**DOI:** 10.1101/2024.11.29.625575

**Authors:** Emily Tolbert, Dalton Dacus, Rose Pollina, Nicholas A. Wallace

## Abstract

Beta genus human papillomaviruses (β-HPVs) are ubiquitous double stranded DNA (dsDNA) viruses that may promote skin cancers by destabilizing the host genome. Supporting this, expression of the E6 gene from a β-HPV (β-HPV 8 E6) results in increased micronuclei that should induce an innate immune response that eliminates these cells. Yet, β-HPV 8 E6 promotes rather than restricts proliferation. We hypothesize that β-HPV 8 E6 accomplishes this by attenuating the cyclic GMP-AMP synthase-stimulator of interferon genes (cGAS-STING) pathway, an innate immune pathway that becomes activated in response to cytosolic micronuclear dsDNA. Here, we show that in response to stimulation by transfection of pLVX-GFP plasmid, β-HPV 8 E6 reduced the magnitude and intensity of cGAS-STING pathway activation in immunoblot experiments. These data also demonstrate that impairment of the cGAS-STING pathway is strongest downstream of STING phosphorylation. Further, RNA-sequencing suggests that β-HPV 8 E6 downregulates other innate immune pathways. We also show that cGAS is recruited to micronuclei induced by β-HPV 8 E6. These data suggest a mechanism by which β-HPV 8 E6 facilitates proliferation of cells destabilized by micronuclei and support the hypothesis that the prevalence of β-HPV infections is in part due to the impairment of the cGAS-STING innate immune response.

## Introduction

Beta human papillomaviruses (β-HPVs) are ubiquitous small, non-enveloped double-stranded DNA (dsDNA) viruses that are commonly spread through skin-to-skin contact (Antonsson et al., 2003). Infections often first occur in early childhood with common reinfections (Weissenborn et al., 2009; Antonsson et al., 2003). While β-HPV infections occur frequently, they are generally transient in nature. Infections typically only last between 8 to 11 months (Hampras et al., 2017). These infections are asymptomatic in immunocompetent people but a subset of β-HPVs, including β-HPV 8, have been shown to increase the risk of skin cancer in people with the rare genetic disorder epidermodysplasia verruciformis and other forms of immunosuppression (Kremsdorf et al., 1982; Pfister et al., 1981; Orth, 1986; Bouwes Bavinck et al., 2001; Hsu et al., 2009). Mechanistic evidence suggests that the E6 protein from these β-HPVs contributes to this increased risk by altering DNA repair mechanisms and creating genomic instability (Wendel & Wallace, 2017). The result of these alterations includes the increased frequency of micronuclei, anaphase bridges, and other mitotic errors (Dacus et al., 2022).

In alpha HPVs, E6 has also been implicated in impairing innate immune responses triggered by pattern recognition receptors (PRRs) in response to pathogen-associated molecular patterns (PAMPs) including the Toll-like receptors (TLRs), RIG-1-like receptors (RLRs), and cyclic GMP-AMP synthase-stimulator of interform genes (cGAS-STING) (Lo Cigno et al., 2020; Albertini et al., 2018; Lau et al., 2015; Luo et al., n.d.; Hasan et al., 2013; Ronco et al., 1998; Okude et al., 2021). The cGAS-STING pathway recognizes dsDNA in the cytosol of cells, such as β-HPVs and micronuclei induced by β-HPV E6 (Sun et al., 2013; Mackenzie et al., 2017). Once cGAS detects and binds to cytoplasmic DNA, it synthesizes the secondary messenger cGAMP (cyclic GMP-AMP). cGAMP then migrates to the endoplasmic reticulum to activate STING (stimulator of interferon genes) (Sun et al., 2013). Once activated, STING translocates to the Golgi apparatus where TBK1 (tank binding kinase) is recruited. TBK1 autophosphorylates itself and phosphorylates STING. Phosphorylated STING then recruits IRF3, a transcription factor, to be phosphorylated by phosphorylated TBK1 (Tanaka & Chen, 2012). Phosphorylated IRF3 translocates to the nucleus to activate the expression of type 1 interferons (Sun et al., 2013). Type 1 interferon induction will trigger the activation of the JAK-STAT pathway which will induce the expression of interferon stimulated genes (ISGs). This expression will culminate in an apoptotic or senescent response (Luo et al., 2022).

While numerous studies have identified an antagonistic relationship between alpha HPVs and PRRs, there are few formal investigations examining the role of β-HPV proteins in these interactions. Here, we use immunoblot and RNA-sequencing approaches to show that β-HPV 8 E6 impairs the cGAS-STING innate immune response. We also show that cGAS-STING signaling factors are recruited to micronuclei induced by β-HPV 8 E6. We provide a detailed molecular analysis of the cGAS-STING perturbation, showing that the impairment begins post-STING phosphorylation. Taken together, these data are consistent with the hypothesis that the prevalence of β-HPV 8 is in part due to the ability of β-HPV8 E6 to attenuate the cGAS-STING pathway.

## Materials and Methods

### Cell Culture

Telomerase immortalized human foreskin keratinocytes (hTERT HFKs) were provided by Michael Underbrink (University of Texas Medical Branch). Cells were grown in Keratinocyte Growth Medium 2 (PromoCell) supplemented with calcium chloride (PromoCell), SupplementMix (PromoCell), and penicillin-streptomycin (Caisson). hTERT HFKs expressed empty vector control (LXSN) and HA-tagged β-HPV8 E6 driven by the promoter in the LXSN vector.

### Transfections

For immunoblots, hTERT HFK cells were plated in 9 mL of complete growth medium in a 10cm plate. Cells were used at 70-80% confluency. In time series transfections, 0.5ug of pLVX-GFP plasmid was diluted in 200 μL Xfect transfection reagent (631317, Takara). In DNA dose response transfections, cells were transfected with selected plasmid concentrations 0ug, 0.25ug, 0.5ug, 1ug, 2.5ug, 5ug diluted in 200 μL Xfect transfection reagent (631317, Takara).

The resulting transfection mixture was incubated at room temperature for 15 min and then the transfection mixture was added to each plate in a drop-wise manner. Time series plates were incubated for selected time intervals (0hrs, 4hrs, 8hrs, 12hrs, 24hrs, 30hrs, and 48hrs) at 37°C. DNA dose response plates were incubated for 20hrs at 37°C. Cells were then harvested for immunoblotting.

For RNA-sequencing, hTERT HFK cells were plated in 20 mL of complete growth medium in a 15 cm plate. Cells were used at 70–80% confluency. Cells were transfected with 1ug of plasmid diluted in 1200 μL Xfect transfection reagent (631317, Takara). The mixture was incubated at room temperature for 15 min. The transfection mixture was added to each well drop-wise and incubated for 20 hours at 37°C. Cells were harvested for RNA extraction and sequencing.

### Immunoblotting

Cells were washed with ice-cold phosphate-buffered saline (PBS) then lysed with radioimmunoprecipitation assay lysis buffer (Santa Cruz Biotechnology) supplemented with phosphatase inhibitor cocktail 2 (Sigma-Aldrich) and protease inhibitor cocktail (Bimake).

Protein concentration was determined with a Pierce bicinchoninic acid (BCA) protein assay kit (Thermo Scientific). Lysates were run on Novex 4 to 12% Tris-glycine WedgeWell mini gels (Invitrogen) and transferred to Immobilon-P membranes (Millipore). Membranes were probed with the following primary antibodies: IRF3 (4302S; Cell Signaling Technology), pIRF3 (Ser396; catalog no. 4947S; Cell Signaling Technology), STING (13647S; Cell Signaling Technology), pSTING (Ser366; catalog no. 19781S; Cell Signaling Technology), TBK1/NAK (3504S; Cell Signaling Technology), pTBK1 (Ser172; 5483S; Cell Signaling Technology), VIPERIN (MABF106; Sigma-Aldrich), and C23 (MS-3) HRP (sc-8031; Santa Cruz Biotechnology). After exposure to the matching appropriate horseradish peroxidase-conjugated secondary antibody, membranes were visualized using SuperSignal West Femto maximum sensitivity substrate (Thermo Scientific). Densitometry quantification was performed using ImageJ (NIH, Rockville, MD, USA).

### Immunofluorescent Microscopy

Cells were seeded into 6 well plates with etched coverslips and grown overnight. Cells were fixed with 4% formaldehyde. Then, 0.1% Triton-X solution in PBS was used to permeabilize the cells, followed by blocking with 3% bovine serum albumin in PBS for 30 min. Cells were then incubated with the following: cGAS (D1D3G; Cell Signaling Technologies), Lamin B1 (C-12; sc-365214; Santa Cruz Biotechnology). The cells were washed and stained with the following appropriate secondary antibodies: Alexa Fluor 594-goat anti-rabbit (Thermo Scientific, catalog no. A11012) and Alexa Fluor 488 goat anti-mouse (Thermo Scientific, catalog no. A11001). After washing, the cells were stained with 2 μM 4′,6-diamidino-2-phenylindole (DAPI) in PBS and visualized with a Zeiss LSM 770 or a Zeiss LSM 880 Airyscan microscope. Laser power, gain, and the pinhole remained the same between experiments to ensure accurate quantification. Background fluorescence was used as an internal control between samples.

Images were analyzed using ImageJ techniques, as described previously (Murthy et al., 2018).

### RNA Isolation and Purification

Cells were washed with phosphate buffered saline then lysed with Trizol reagent. RNA was then isolated using Zymo Clean and Concentrator Kit (Zymo Research). Sample purity was assessed using nanodrop software. Samples were then delivered to Molecular and Cell Biology Core (NIH COBRE, Center on Emerging and Zoonotic Infectious Diseases) at Kansas State University for sequencing.

### RNA-Sequencing

Two microliters of each sample were used for quantification using Qubit™ RNA BR Assay Kit (Thermo Fisher Scientific) according to manufacturer’s instructions in the Qubit fluorometer 4 (Thermo Fischer Scientific). After quantification all samples were normalize to 40 ng/µl (total 1000 ng). The samples were also analyzed using Tapestation 4150 (Agilent) using RNA ScreenTape Analysis (Agilent) following manufacturer’s instructions. All samples had RIN (RNA Integrity Number) above 9.4. The library preparation was performed using the kit Illumina Stranded mRNA Prep (Illumina) according to manufacturer’s instructions. Briefly, the messenger RNAs (mRNAs) are purified using oligo(dT) magnetic beads. Then, the mRNAs are used as template to synthesize the first strand cDNA followed by the synthesis of the second strand cDNA. After the double strand cDNA synthesis, the samples were submitted to a reaction to add adenine nucleotide to the 3’ ends, here called anchors, to the blunt fragments to prevent the fragments from ligate to each other during the adaptor ligation step. The next step was the library amplification, which specifically amplifies anchor-ligated DNA and adds the indexes and primer sequences for clustering generation. For this library prep the index adapters used was IDT-ILMN Nextera DNA UD Indexes Set A. After the amplification step, the library was cleaned up using AMPure XP (Beckman Coulter), according to manufacturer’s protocol. The samples were quantified using Qubit dsDNA HS Assay Kit (Thermo Fisher Scientific) according to manufacturer’s instructions in the Qubit fluorometer 4 (Thermo Fischer Scientific) (Table 3). The samples were also evaluated on Tapestation 4150 (Agilent) using the kit High Sensitivity D5000 (Agilent) and the average size of the libraries was 284.25. The samples were dilute to 4 nM and 10µl of each sample were pooled into a 1.5 ml low-binding tube labeled “library pool”. For the denaturation and dilution, the “Standard Normalization Method” from “NextSeq Denature and Dilute Libraries Guide was performed. Briefly, 0.2 N NaOH was used to denature the 4 nM library pool followed by dilution with HT1 buffer until the concentration of 1.8 pM (loading concentration). 1% of PhiX (Illumina) was used as run control. The run was performed using NextSeq 500/550 High Output Kit v2.5 (150 Cycles) reagent kit (Illumina) in the NextSeq 550 sequencing system (Illumina).

### RNA-Sequencing Analysis

Data quality was assessed using FASTQC analysis (Andrews, 2010). Paired end reads were then aligned using a custom hg38 human reference genome appended with β-HPV 8 E6 gene with STAR aligner (version 2.7.10b) (Dobin et al., 2013). Once aligned, gene counts were generated using FeatureCounts (Liao et al., 2014). Differential expression analysis was performed in R studio using DESeq2 (Love et al., 2014). DEGs were identified by applying a p-value < 0.05 cutoff. These results were used to generate volcano plots in R studio using ggplot2 package. Gene ontology was performed using PANTHERGO (Mi et al., 2019). Innate immunity associated genes were identified using InnateDB (Lynn et al., 2008).

### Statistical Analysis

Statistical significance was determined using student’s t-test. Results were reported as significant for all p values less than 0.05.

## Results

### E6 reduces cGAS-STING signaling in response to cytosolic DNA

We hypothesized that that β-HPV 8 E6 hindered activation of the cGAS-STING pathway.

To examine this hypothesis, we transfected cells with pLVX-GFP plasmid (500ng) to stimulate the cGAS-STING pathway and collected protein lysates at intervals for 48 hours. We then used immunoblots to detect the abundance of cGAS-STING proteins in vector control keratinocytes (hTERT HFK LXSN) and keratinocytes expressing β-HPV 8 E6 (hTERT HFK E6). The pLVX-GFP plasmid expresses green fluorescent protein (GFP), allowing us to confirm transfection success using fluorescence microscopy. In hTERT HFK LXSN cells, cGAS, IRF3 (total and phosphorylated), STING (total and phosphorylated) and a representative ISG (viperin) were all stimulated by transfection with pLVX-GFP (**Fig 1**). The duration and magnitude of multiple indicators of cGAS-STING were comparatively decreased in hTERT HFK E6 cells (**Fig 1**).

**Figure 1.**
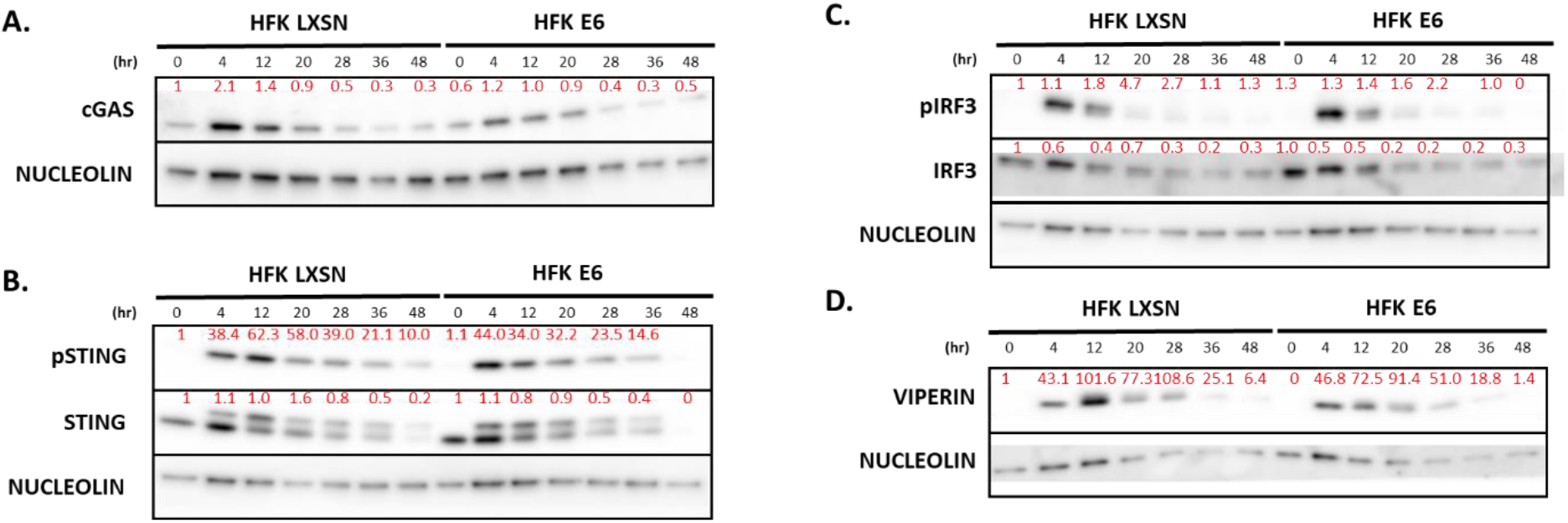
E6 truncates the response of cGAS-STING signaling. Representative immunoblots of time-course transfected hTERT HFK LXSN (HFK LXSN) and hTERT HFK 8 E6 (HFK E6) cell lines. NUCLEOLIN (C23) was used as a loading control for all immunoblots. Cells were transfected with 0.5ug of pLVX-GFP plasmid then harvested 0-48 hours post transfection. Immunoblot experiments were repeated in n=2. Red numbers above bands show densitometry quantification normalized to HFK LXSN 0 hour treatment and NUCLEOLIN loading control. (A) Representative immunoblot of cGAS abundance. (B) Representative immunoblot of STING (total and phosphorylated) abundance. (C) Representative immunoblot of IRF3 (total and phosphorylated) abundance. (D) Representative immunoblot of VIPERIN abundance.

Our time course analysis indicates that the largest differences between hTERT HFK LXSN and hTERT HFK E6 cells occurred 20 hours after transfection. Thus, to further characterize the ability of E6 to suppress cGAS-STING pathway activation, we transfected hTERT HFK LXSN and hTERT HFK E6 cells with a gradient of pLVX-GFP plasmid concentrations (0ug-5ug) and harvested protein lysates at 20 hours post transfection.

Immunoblot analysis found that these transfections resulted in reproducible activation of the cGAS-STING pathway in hTERT HFK LXSN (**Fig 2**). However, these responses were more muted in hTERT HFK E6 cells.

**Figure 2.**
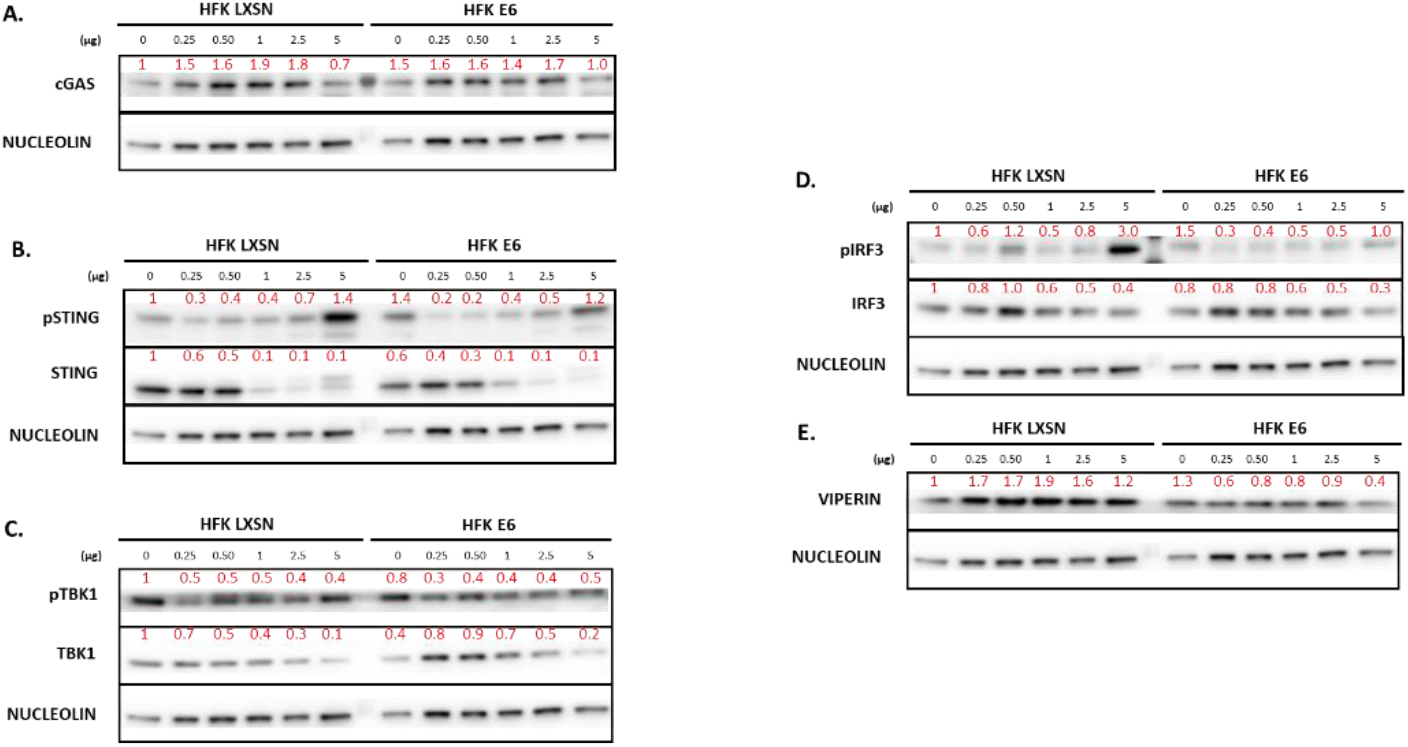
E6 decreases abundance of cGAS-STING pathway in response to dose gradient. Representative immunoblots of dsDNA-dose response transfected hTERT HFK LXSN (HFK LXSN) and hTERT HFK 8 E6 (HFK E6) cell lines. Immunoblot experiments were repeated in n=2. NUCLEOLIN (C23) was used as a loading control for all immunoblots. Cells were transfected with 0-5ug of pLVX-GFP plasmid then harvested 20 hours post transfection. Red numbers above bands show densitometry quantification normalized to HFK LXSN 0ug treatment and NUCLEOLIN loading control. (A) Representative immunoblot of cGAS abundance. (B) Representative immunoblot of STING (total and phosphorylated) abundance. (C) Representative immunoblot of TBK1 (total and phosphorylated) abundance (D) Representative immunoblot of IRF3 (total and phosphorylated) abundance. (E) Representative immunoblot of VIPERIN abundance.

We previously reported that hTERT HFK E6 cells have more micronuclei than hTERT HFK LXSN cells (Dacus et al., 2022). Because cGAS-STING signaling is also activated in response to micronuclei, we analyzed cGAS-STING activation in the micronuclei of hTERT HFK LXSN and hTERT HFK E6 cells (**Fig 3**). Immunofluorescent microscopy can detect cGAS-STING activation by localization of cGAS to micronuclei (Harding et al., 2017; Mackenzie et al., 2017). There was no significant difference in cGAS localization to micronuclei in hTERT HFK E6 cells.

**Figure 3.**
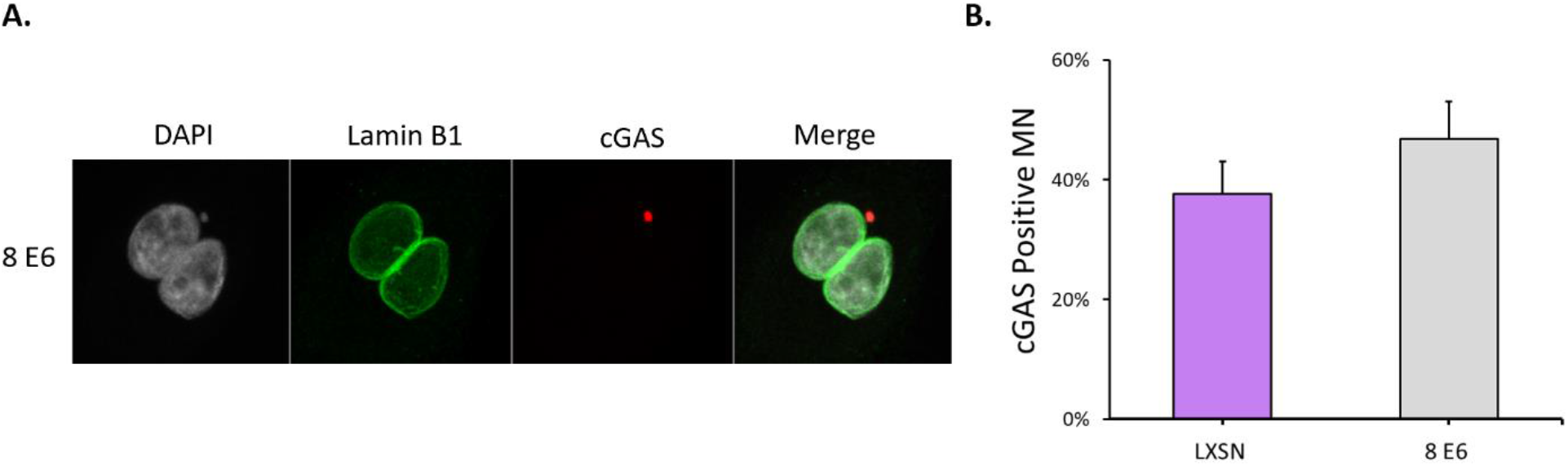
Functional cGAS detection in hTERT HFK E6 cells. Representative immunofluorescent images of cGAS localization to micronuclei in hTERT HFK E6 cells. (B) Quantification of cGAS positive micronuclei in hTERT HFK LXSN and hTERT HFK E6 cells. Results were averaged between three independent experiments. At least 250 micronuclei/cell line were quantified for cGAS frequency across the three experiments. Graph depicts ± the standard errors of the mean.

### E6 impairs innate immune responses

Our immunoblot data indicates that β-HPV8 E6 decreases cGAS-STING signaling in response to cytoplasmic DNA. To more broadly characterize the functional effectiveness of the innate immune system, we performed Illumina RNA-sequencing on hTERT HFK LXSN and hTERT HFK E6 cells 20 hours after transfection with 1ug of pLVX-GFP plasmid. These conditions were chosen because they maximized the differences in cGAS-STING signaling between hTERT HFK LXSN and hTERT HFK E6 cells. Sequencing was performed using Illumina NextSeq 550 sequencing system. After sequencing, data quality was analyzed using FASTQC. Samples were then aligned using STAR aligner and gene counts were quantified using FeatureCounts. Counts were then used to perform differential gene expression (DEG) analysis using DESeq2. When comparing hTERT HFK LXSN untreated and hTERT HFK LXSN treated samples, 622 DEGs were identified. Of these DEGs, 453 were upregulated and 169 were downregulated (**Fig 4A)**. For hTERT HFK LXSN cells, upregulated genes were significantly correlated with innate immune activation, as denoted by triangles in **Fig 4A**. When comparing hTERT HFK E6 treated and untreated cells, there were 373 DEGs. Of these, 15 were upregulated and 358 were downregulated (**Fig 4B**). No upregulated genes were identified as being associated with innate immunity, while 25 downregulated genes were identified as innate immune genes.

**Figure 4.**
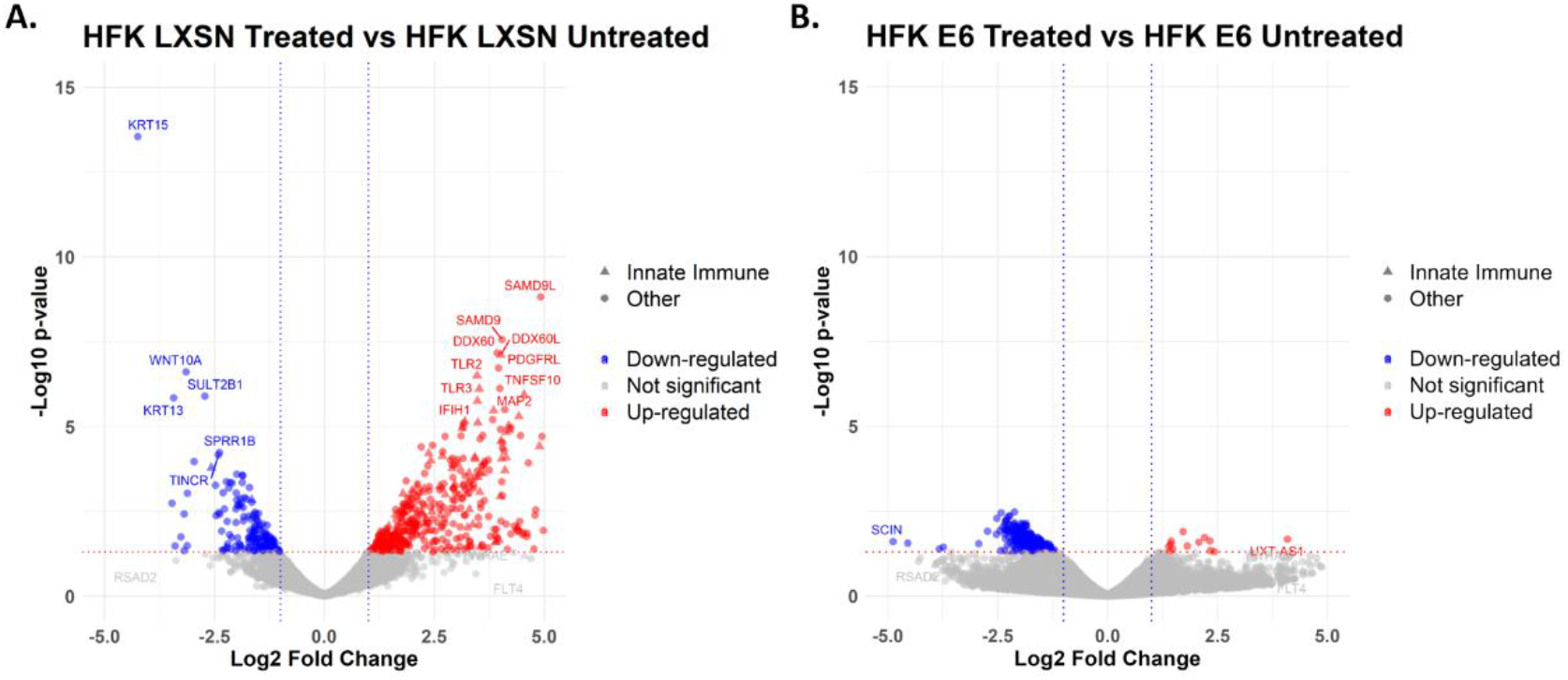
E6 impairs innate immune responses. hTERT HFK LXSN (HFK LXSN) and hTERT HFK E6 (HFK E6) cells were transfected with 1ug of pLVX-GFP plasmid and harvested 20 hours post-transfection. RNA was isolated and purified then submitted for Illumina sequencing. Differentially expressed genes were identified using DESeq2 software. Significant differentially expressed genes were identified as having p-values less than 0.05. (A) Volcano plot of significant differentially expressed genes between hTERT HFK LXSN treated and hTERT HFK LXSN untreated cells. (B) Volcano plot of significant differentially expressed genes between hTERT HFK E6 treated and hTERT HFK E6 untreated cells.

## Discussion

The ubiquity of β-HPVs suggests that they possess a mechanism for evading immune responses. The cGAS-STING innate immune response should effectively suppress β-HPV infections and prevent the genome-destabilizing mutations that arise because of infections.

While we have shown that cGAS is capable of detecting dsDNA in the form of micronuclei in the cytoplasm of immortalized keratinocyte expressing β-HPV 8 E6 and vector control LXSN cells (**Fig 3**), downstream pathway analysis revealed significant impairment of the pathway in hTERT HFK E6 cells. Pathway activation was attenuated in hTERT HFK E6 cells before the 48-hour timepoint (**Fig 1**) and key pathway components displayed decreased abundance in immunoblot experiments (**Fig 2**). These changes seem to occur post-STING phosphorylation as seen in **Figure 5** in hTERT HFK E6 cells. These findings build on the observations noted in Rattay et. al (2022) that β-HPV 8 E6 does not impair STING expression at a transcriptional level and instead impacts cGAS-STING signaling at a post translation modification level. A functional innate immune response was detected in hTERT HFK LXSN cells through gene ontology analysis with both ISGs and IFNs upregulated. In hTERT HFK E6 cells, there are no upregulated innate immune associated genes as identified by gene ontology. There are also a number of downregulated innate immune genes associated with additional immune signaling pathways, most notably, RIG-1 and TLR signaling pathways. These results suggest that E6 has a broad ability to suppress innate immune responses but does not fully attenuate innate immune signaling.

**Figure 5.**
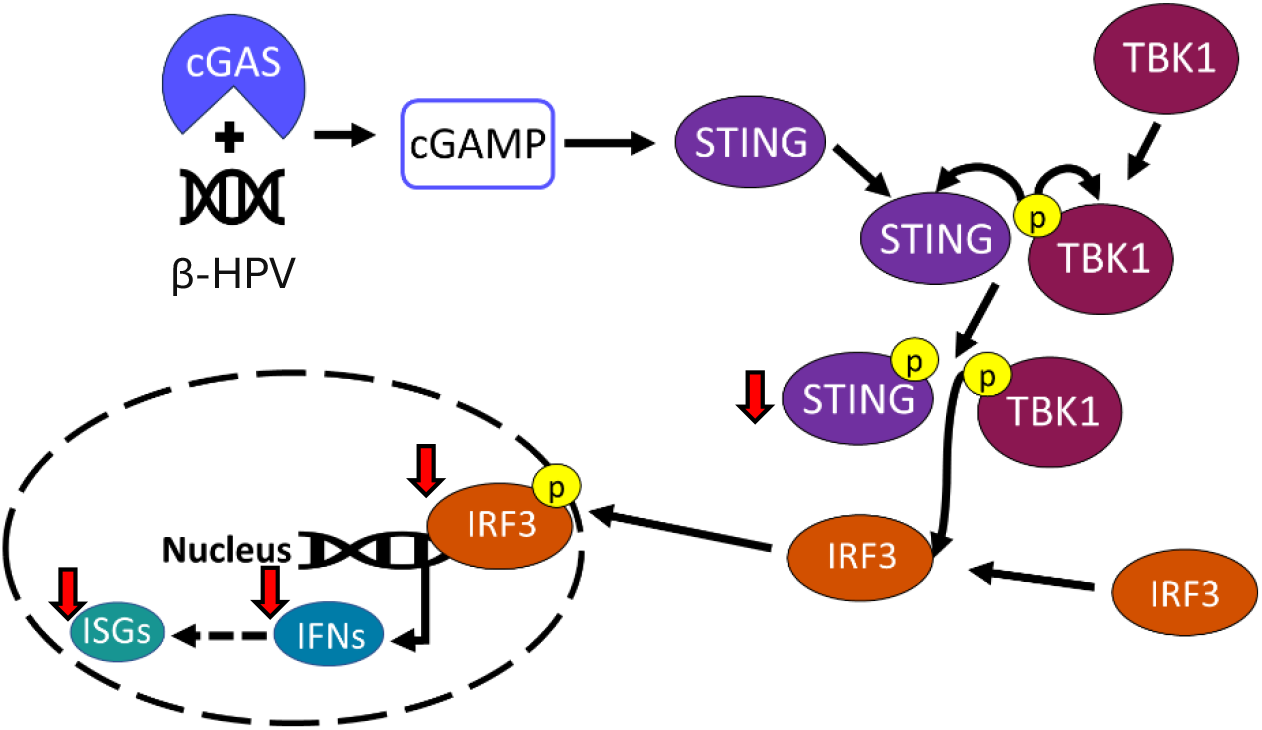
Summary Figure of Changes Induced by Beta HPV 8 E6. Red arrows denote proteins that display decreased abundance in immunoblotting experiments or downregulation in RNA-sequencing experiments.

β-HPVs face a selective pressure to evolve mechanisms that evade or suppress immune responses, thereby enhancing their ability to propagate infections. This evolutionary process can also lead to an increased potential to cause genomic instability by altering cell signaling pathways (Wallace et al., 2014; Dacus et al., 2020). One consequence of this is micronuclei (Ye et al., 2019). If cells containing micronuclei are not properly targeted for apoptosis, the micronucleus can reintegrate into host nuclei and potentially cause chromothripsis, which is a highly mutagenic event (Zhang et al., 2015). Functioning innate immunity is therefore critical to safeguarding genome stability.

While this work highlights the ability of β-HPV 8 E6 to impair the cGAS-STING innate immune response, there are still questions as to what extent β-HPVs impair innate immunity. While our work focuses on the E6 protein, it is unclear what effect the expression of the complete early region would have on cGAS-STING signaling. Rattay et. al (2022) identified roles for β-HPV 8 E1 and E2 genes to antagonize PRRs in cells. There are also several documented roles for E7 in impairing cGAS-STING signaling (Lou et al., 2023; Lo Cigno et al., 2020). We have also yet to determine a mechanism for pathway impairment post STING phosphorylation. Finally, while other papers have examined β-HPV effect on RIG-1 and TLR signaling (Chiang et al., 2018; Rattay et. al, 2022), the full impairment of the innate immune response has yet to be characterized at a transcriptional and post-translational modification level.

